# Repeat-rich regions cause false positive detection of NUMTs: a case study in amphibians using an improved cane toad reference genome

**DOI:** 10.1101/2024.07.04.601973

**Authors:** K Cheung, LA Rollins, JM Hammond, K Barton, JM Ferguson, HJF Eyck, R Shine, RJ Edwards

## Abstract

Mitochondrial DNA (mtDNA) has been widely used in genetics research for decades. Contamination from nuclear DNA of mitochondrial origin (NUMT) can confound studies of phylogenetic relationships and mtDNA heteroplasmy. Homology searches with mtDNA are widely used to detect NUMTs in the nuclear genome. Nevertheless, false positive detection of NUMTs is common when handling repeat-rich sequences, whilst fragmented genomes might result in missing true NUMTs. In this study, we investigated different NUMT detection methods and how the quality of the genome assembly affects them. We presented an improved nuclear genome assembly (aRhiMar1.3) of the invasive cane toad (*Rhinella marina*) with additional long-read Nanopore and 10x linked-read sequencing. The final assembly was 3.47 Gb in length with 91.3% of tetrapod universal single-copy orthologs (n=5,310), indicating the gene-containing regions were well assembled. We used three complementary methods (NUMTFinder, *dinumt* and *PALMER*) to study the NUMT landscape of the cane toad genome. All three methods yielded consistent results, showing very few NUMTs in the cane toad genome. Furthermore, we expanded NUMT detection analyses to other amphibians and confirmed a weak relationship between genome size and the number of NUMTs present in the nuclear genome. Amphibians are repeat-rich, and we show that the number of NUMTs found in highly repetitive genomes is prone to inflation when using homology-based detection without filters. Together, this study provides an exemplar of how to robustly identify NUMTs in complex genomes when confounding effects on mtDNA analyses are a concern.

**Significance:** This study uses an updated cane toad nuclear genome assembly and multiple NUMT detection methods to confirm a lack of NUMTs that might confound the use of mtDNA as a population genetic marker in the cane toad. We provide an exemplar study for NUMT detection accounting for genome assembly quality and composition, and highlight the risks of using BLASTN-based approaches in highly repetitive nuclear genomes.

## Introduction

Mitochondrial DNA (mtDNA) has been utilised in phylogenetics, DNA barcoding and population genetics studies for decades. High copy numbers per cell make mitochondrial DNA easier to retrieve from samples of low quantity or quality. Additionally, the haplotype nature, mode of inheritance, and abundant knowledge of mitochondrial genomes contribute to their usefulness in genetics research. Studies of mitochondrial DNA have yielded rich bodies of research on the evolution, population structure, and phylogenetics of a broad range of taxa (Ballard and Rand 2005; Gray 2012; Kivisild 2015).

One issue that requires consideration when analysing mtDNA data is the presence of nuclear DNA of mitochondrial origin (NUMTs). Numtogenesis is the transferal of mtDNA into the nuclear genome, creating NUMTs (Singh, et al. 2017). NUMTs can become integrated into the nuclear genome when double-strand breaks are repaired by non-homologous end joining (NHEJ) (Hazkani-Covo and Covo 2008) or microhomology-mediated end joining (Wei, et al. 2022). A large-scale human study of tumour and germline cells revealed an enrichment of NUMTs in the non-coding D-loop in mitochondrial genomes from tumour cells, in relation to the transcription start site and origin of replication (Wei, et al. 2022). NUMTs have been reported across almost every taxonomic group in eukaryotes and a positive relationship between the nuclear genome size and total NUMT content in the genome has been reported (Bensasson, et al. 2001; Hazkani-Covo, et al. 2010; Richly and Leister 2004).

However, this relationship is not universal. For example, the Zebrafish nuclear genome (1.4 - 1.7 Gb in length) does not contain NUMTs (Hazkani-Covo, et al. 2010) whilst for avian species with similarly small genomes (0.91 – 1.3 Gb) (Zhang, et al. 2014), the abundance of NUMTs varies (ranging from 4 to > 600 NUMTs) (Liang, et al. 2018). The variation in abundance of NUMTs across different taxonomic groups could result from differences in the rate of NUMT insertion, the rate of NUMT duplication and the rate of NUMT removal (Hazkani-Covo, et al. 2010).

NUMT sequences are expected to diverge from their mtDNA counterparts over time, as the nuclear copy will not be under functional constraint. However, at their creation, NUMTs are exact copies of their mitochondrial sequence of origin and can remain similar for extended periods of time due to the low mutation rate of nuclear DNA. As a consequence, NUMTs may be amplified by mtDNA PCR primers, leading to incorrect interpretation of data.

Because the age of NUMTs is unknown *a priori*, it is important to investigate the potential for NUMT contamination of mtDNA data. When undetected, NUMTs can cause inaccuracies in estimates derived from mtDNA data. For example, NUMTs can lead to an overestimation of the number of novel mutations on a population scale, potentially resulting in inaccurate inferences related to allelic frequency, population structure, demography, and phylogenetic relationships, or misidentification of heteroplasmy (Maude, et al. 2019; Schultz and Hebert 2022; Song, et al. 2008). Methods to detect NUMTs include bioinformatic prediction (Dayama, et al. 2014; Dayama, et al. 2020; Hebert, et al. 2023; Zhou, et al. 2020) and laboratory-based detection (PCR amplifications) (Kuprina, et al. 2023; Machida and Lin 2017), but their implementation is often restricted by cost and efficiency.

Previously, we investigated mitochondrial population genomics of the notorious cane toad (*Rhinella marina*) invasion from its native range in French Guiana to Hawai’i, and subsequently to Australia (Cheung, et al. 2024). This invasion is well-known for a broad collection of evidence of rapid evolution following introduction (Rollins, et al. 2015). Although cane toads were only introduced to Australia in the 1930’s, we detected a large number of mitochondrial genetic variants private to the Australian introduced range (Cheung, et al. 2024). The haplotype network formed a star-shaped topology, indicating that these variants arose recently. Such a result could indeed arise from the incorporation of NUMTs, but our explicit search for mitochondrial insertions in the nuclear genome did not identify any (Cheung, et al. 2024). However, because genome quality is known to be important to the detection of NUMTs (Triant and Pearson 2022), that study may have been limited by the use of a relatively incomplete draft nuclear genome, hereon refered to as aRhiMar1.2 (Edwards, et al. 2018). Furthermore, only a single approach was used to detect NUMTs (Edwards, et al. 2021), which uses BLASTN to query the nuclear genome for mitochondrial sequences. Although the use of homology searching is the most widely used approach to detect NUMTs (e.g. (Hazkani-Covo, et al. 2010; Hebert, et al. 2023; Richly and Leister 2004)), it has several pitfalls. This method is susceptible to false positives arising from random alignments, especially when dealing with sequences rich in repeats, even when stringent E-values are applied. Moreover, in cases where the genome assembly is highly fragmented, NUMTs might have failed to assemble correctly and searches may fail to detect NUMTs located at assembly gaps. An alternative approach involves using raw short-read or long-read sequencing data for the detection process. Whilst harder to implement, this approach offers an advantage because it directly identifies NUMTs within raw reads, bypassing genome quality and/or alignment issues. For example, *dinumt* (Dayama, et al. 2014) utilises paired-end short-reads in which the split read pairs map to the nuclear genome and mitochondrial genome respectively. *dinumt* identifies candidate NUMTs by clustering the mapped paired reads based on the mapping positions and orientation. *PALMER* (Zhou, et al. 2020) uses long-reads to detect non-reference mobile element insertions (MEI) and with modification, can be expanded to detect other categories of non-reference insertion (e.g. NUMTs). Combining several of these methods could help to reduce false-positives from homology methods and false-negatives from reads-based methods, potentially leading to more accurate detection of NUMTs than any single approach.

In this study, we aimed to clarify the NUMT landscape in the cane toad genome. Specifically, we significantly improved the quality of the cane toad nuclear genome (aRhiMar1.3) to examine how this affects the detection of NUMTs. Additionally, we compared two alternative NUMT detection methods (*dinumt* and *PALMER*) with NUMTFinder. Because little is known about NUMTs in amphibians, we also assessed whether the lack of NUMTs we found in the cane toad genome is representative of other amphibians. This robust approach to NUMT detection furthers our understanding of evolutionary processes in the introduced cane toad population in Australia but also provides a model for other mitogenome studies aiming to remove the potential confound of NUMT inclusion.

## Materials and Methods

### DNA sequencing

Raw Illumina 2 x 150 bp paired end and PacBio Continuous Long Read genomic sequencing data from the original cane toad genome (Edwards, et al. 2018) were downloaded from National Centre for Biotechnology Information (NCBI) BioProject PRJEB24695. The same high-molecular-weight genomic DNA (gDNA) (see (Edwards, et al. 2018) for details) was used for additional 10x linked-read and Oxford Nanopore Technologies (ONT) sequencing. A 10x Genomics (Pleasanton, USA) linked-read library was prepared using the Chromium™ Genome Reagent Kit (v2 Chemistry) and sequenced (2 × 150 bp paired-end) on the Illumina HiSeq X Ten platform. For ONT sequencing, a combination of five libraries were sequenced on PromethION (one) and GridION (four). For PromethION sequencing, 1 μg gDNA was prepared using the Genomic DNA by ligation protocol (SQK-LSK109) according to the manufacturer’s instructions. The final library prep was loaded onto a FLO-PRO002 PromethION flow cell (pore version 9.4.1). For GridION sequencing, four libraries were made by preparing 400 ng gDNA using the rapid barcoding protocol (SQK-RBK004) according to the standard protocol. Each library was loaded onto a FLO-MIN106 GridION flow cell (R9.4.1 chemistry). The sequencing was run for 72 hours. GPU-enabled guppy (v3.3.0) (Wick, et al. 2019) base-calling was performed after sequencing (PromethION high accuracy flip-flop model; config ‘dna_r9.4.1_450bps_hac.cfg’ config). The resulting FASTQ files were combined and filtered with custom script.

### Genome assembly

The assembly workflow involved assembling a draft long-read assembly, hybrid polishing with long- and short-reads and gap-filling with long-reads. Computational tasks were carried out on the computational cluster Katana supported by Research Technology Services at UNSW Sydney (PVC Research Infrastructure).

Due to the discrepancy in the genome size estimation among cell-based (flow cytometry: 3.98 to 4.90 Gb (Chipman, et al. 2001; MacCulloch, et al. 1996); densitometry: 4.06 to 5.65 Gb (Bachmann 1970; Goin, et al. 1968)), *k*-mer based and qPCR approaches (1.77 to 2.3 Gb; 2.38 Gb (Edwards, et al. 2018)), we employed an alternative method, DepthSizer v1.8.0 (Chen, et al. 2022b), utilising long-read sequencing depth profiles of single-copy BUSCO genes to estimate the genome size. This estimated the size at approximately 3.5 Gbp, which was used for guiding the subsequent assembly.

The first step was to use Flye v2.7b (Kolmogorov, et al. 2019) to assemble both PacBio long-reads and Nanopore long-reads. The initial assembly was tidied with Diploidocus v0.15.0 (Chen, et al. 2022b) (default dipcycle mode). Hybrid polishing was performed using HyPo v1.0.3 (Ritu, et al. 2019) with with 10x Chromium reads and mixed long-read data mapped onto the genome using LongRanger v2.2.2 (Marks, et al. 2019) and Minimap2 v2.17 (Li 2018), respectively. The HyPo-polished assembly was scaffolded with 10x Chromium reads using ARCS v1.1.1 (Yeo, et al. 2018) and gap-filled with both PacBio long-reads and Nanopore long-reads using SSPACE-LongRead (March 2021) (Boetzer and Pirovano 2014) and GapFinisher v20190917 (Kammonen, et al. 2019). The assembly was corrected with a second-round of HyPo hybrid polishing, followed by a final purging of false duplicates and low quality sequences with Diploidocus. Contigs corresponding to pure mtDNA were identified with NUMTFinder v0.5.1 (Edwards, et al. 2021) and removed from the assembly to produce the final nuclear reference genome. The published high-quality mtDNA reference genome (Cheung, et al. 2024) was then added back to the genome (aRhiMar1.3).

### Genome annotation and evaluation

The genome was annotated using the homology-based gene prediction program GeMoMa v1.7beta (Keilwagen, et al. 2019) and 10 protein annotations from 10 species (*Anolis carolinensis* (GCA_000090745.2 (Alfoldi, et al. 2011))*, Canis lupus familiaris* (GCA_014441545.1)*, Danio rerio* (GCA_000002035.4 (Howe, et al. 2013))*, Gallus gallus* (GCA_016699485.1)*, Homo sapiens* (GCA_000001405.29)*, Mus musculus* (GCA_000001635.9 (Church, et al. 2011))*, Naja naja* (GCA_009733165.1 (Suryamohan, et al. 2020))*, Taeniopygia guttata* (GCA_003957565.2 (Rhie, et al. 2021))*, Takifugu rubripes* (GCA_901000725.2)*, Xenopus tropicalis* (GCA_000004195.4 (Hellsten, et al. 2010))) available on Ensembl release 103 (Yates, et al. 2020). The GeMoMaPipeline function was executed with a maximum intron size of 200kb and “`GeMoMa.Score=ReAlign AnnotationFinalizer.r=SIMPLE AnnotationFinalizer.p=RMARP pc=true o=true`” parameters. Longest isoform per gene were extracted from the annotation and assessed using BUSCO v5.4.3 in proteome mode.

Transposable elements (TEs) were discovered using RECON, RepeatScout and LtrHarvest implemented in RepeatModeler2.0.3 (Flynn, et al. 2020) to build a de novo repeat library. TEs and repetitive elements were searched using the de novo repeat library using RepeatMasker v4.1.2 with RMBlast as the search engine for protein-coding gene annotation. The annotation table was made using the buildSummary.pl script from RepeatMasker. The tRNAs were annotated using tRNAScan v2.0.11 (Chan, et al. 2021). The script “EukHighConfidenceFilter” in tRNAScan was used to filter low-confidence tRNAs.

We assessed the completeness of the genome assembly using Benchmarking Universal Single-Copy Orthologs (BUSCO) v5.4.3 (Manni, et al. 2021), implementing BLAST+ v2.11.0, HMMer v3.3.2, Metaeuk v 6-a5d39d9 and SEPP v4.5.1 with genome mode and the lineage “tetrapoda_odb10” set of 5310 core genes. Assembly quality (QV) was estimated using *k*- mer analysis of 10X reads by Merqury v1.3 (Rhie, et al. 2020). We also employed DepthKopy v1.3.0 (Chen, et al. 2022b) to identify possible over-assembly (collapsed regions) and under-assembly (false duplications or low-quality regions) in the genome assembly.

“Complete” BUSCO genes were compiled across the cane toad and 16 other amphibian genomes (*Bombina bombina*, *Bufo bufo*, *Bufo gargarizans*, *Dendropsophus ebraccatus, Discoglossus pictus, Geotrypetes seraphini*, *Hymenochirus boettgeri*, *Leptobrachium ailaonicum*, *Leptobrachium leishanense*, *Microcaecilia unicolor*, *Pyxicephalus adspersus*, *Rana temporaria*, *Rhinatrema bivittatum*, *Xenopus borealis, Xenopus laevis*, *Xenopus tropicalis*) (Table 4).

### NUMT detection and validation in the cane toad nuclear genome

NUMT detection was compared between aRhiMar1.2 (Edwards, et al. 2018) and aRhiMar1.3 to determine whether genome assembly quality affects the detection of NUMTs. NUMTFinder (Edwards, et al. 2021) is an open-source tool to identify putative NUMTs through BLASTN and collapse nearby NUMT fragments into NUMT blocks. We constructed species-specific repeat library using RepeatModeler (http://www.repeatmasker.org/RepeatModeler/) and masked aRhiMar1.3 using RepeatMasker with default parameters (Tarailo-Graovac and Chen 2009). We used NUMTFinder v0.5.4 to search for fragments of the high-quality mitogenome in both full and masked versions of aRhiMar1.3 as described in Cheung, et al. (2024). We used *dinumt* package (Dayama, et al. 2014) to identify NUMTs with the customised setup “`-- include_mask --ensembl --mask_filename =refNUMTs.bed`”. For long-read sequencing reads, we utilised reads from both PacBio and Nanopore technologies, generated for the cane toad genome assembly. We adapted the package *PALMER* (Zhou, et al. 2020) which detects non-reference mobile element insertion events with long-read sequencing data. The detection could be expanded to the detection of the NUMT insertion with a customised setup “`--ref_ver other --type CUSTOMIZED –-mode raw --TSD_finding FALSE --custom_seq custom_seq.fasta`”.

To validate the putative NUMTs insertions found by raw sequences based approaches, we manually inspected the reads-to-genome alignment files (BAM files) using Integrative Genomic Viewer (IGV) (Robinson, et al. 2011; Thorvaldsdottir, et al. 2013). Only insertions supported by at least 10% of the sequencing depth (PacBio: 22X, 3+ reads; ONT: 12.5X, 2+ reads) were considered genuine insertions to minimise false-positive detection.

### NUMTs detection in amphibian species

To investigate whether the low number of NUMTs in the cane toad genome is a species-specific event or a common occurrence across the family, we analysed a diverse range of amphibian species with chromosome-level genome assemblies and complete mitochondrial genomes available in NCBI (assessed on: 2023-02-01) (Table 4). For each species, we obtained both the nuclear genome and mitochondrial genome using the NCBI Datasets command line tools. Given the significant computational resources required for applying both *dinumt* and *PALMER* to the entire dataset of amphibian raw sequencing data, we opted for the NUMTFinder approach for this analysis. The relationship of genome size and NUMT count was statistically tested with Wilcoxon test.

## Results

### Updated cane toad nuclear genome assembly

We assembled an updated cane toad genome (aRhiMar1.3) using a combination of newly generated Nanopore (ONT) long reads and 10x Genomics linked-reads, along with the previously published PacBio CLR and Illumina PE data from the same individual. We generated 7,245,603 ONT long reads with 44 Gb of sequences, averaging 5.7 kb in length and N50 read length of 11,549 bp. Only reads longer than 500 bps were retained, resulting in 43.7 Gb of sequences. These were co-assembled with the original PacBio data using Flye (Kolmogorov, et al. 2019), yielding an initial assembly of 34,696 contigs with contig N50 of 451 kb and BUSCO completeness at 80.3%. Additionally, we generated 1,179,094,866 paired 10x linked-reads with 349.41 Gb of sequences for scaffolding and error correction. Multiple rounds of polishing, scaffolding, gap-filling and tidying (Supplementary Tables 1) generated the final assembled genome with a length of 3.47 Gb on 7,124 scaffolds (13,172 contigs) (Table 1). The final contig N50 was 859 kb, a 5-fold increase compared with the previous draft genome. The scaffold N50 reached 2.5 Mb. DepthSizer analysis of both versions of the genome assembly estimated that the genome size of the cane toad is 3.4 Gb to 3.5 Gb (IndelRatio mode), consistent with the final assembly size.

**Table 1:** Genome assembly statistics and the BUSCO score of the draft genome assembly (aRhiMar1.2) and the updated *Rhinella marina* assembly, aRhiMar1.3. BUSCO scores were shown in the following notation: (C:complete [S:single, D:duplicated], F:fragmented, M:missing, n:gene number).

|  | **aRhiMar1.2** | **aRhiMar1.3\*** |
| --- | --- | --- |
| Total length (bp) | 2,551,760,159 | 3,473,313,001 |
| Number of scaffolds | NA | 7,124 |
| Longest scaffold (bp) | NA | 15,571,941 |
| Mean scaffold length (bp) | 81,287 | 487,551 |
| Median scaffold length (bp) | 38,590 | 30,657 |
| Scaffold N50 (bp) | NA | 2,502,259 |
| Scaffold L50 | NA | 377 |
| Number of contigs | 31,192 | 13,172 |
| Contig N50 (bp) | 167,498 | 859,429 |
| Contig L50 | 3,373 | 1,052 |
| Gap (N) length (bp) |  | 8,571,305 (0.25%) |
| GC content (%) | 43.23 | 43.75 |
| **BUSCO completeness** |  |  |
| Lineage:Tetrapoda<br>(Genome mode) | C:85.8% [S:83.7%, D:2.1%],<br>F:4.7%, M: 9.5%, n=5310 | C:91.3% [S:89.6%, D:1.7%],<br>F:2.7%, M:6.0%, n=5310 |
| Lineage: Tetrapoda<br>(Proteome mode) |  | C:93.6% [S:91.5%, D:2.1%],<br>F:1.8%, M:4.6% |

\*Statistics included mtDNA.

The quality of the final assembly was assessed with different methods. BUSCO evaluation against the tetrapoda_odb10 dataset (*n*=5310) revealed that aRhiMar1.3 assembly included 91.3% of the complete conserved single-copy genes (Figure 1). This is comparable to the 16 other amphibian genomes analysed, with only *Xenopus* species having notably fewer missing BUSCO genes (Supplementary Figure 1). The completeness and quality value (QV) from Merqury analyses of the cane toad genome assembly yielded 95.4 and 32.4 respectively, indicating high completeness and over 99.99% accuracy in nucleotide-level (error rate = 5.81e^-4^) of the genome assembly.

**Figure 1:**
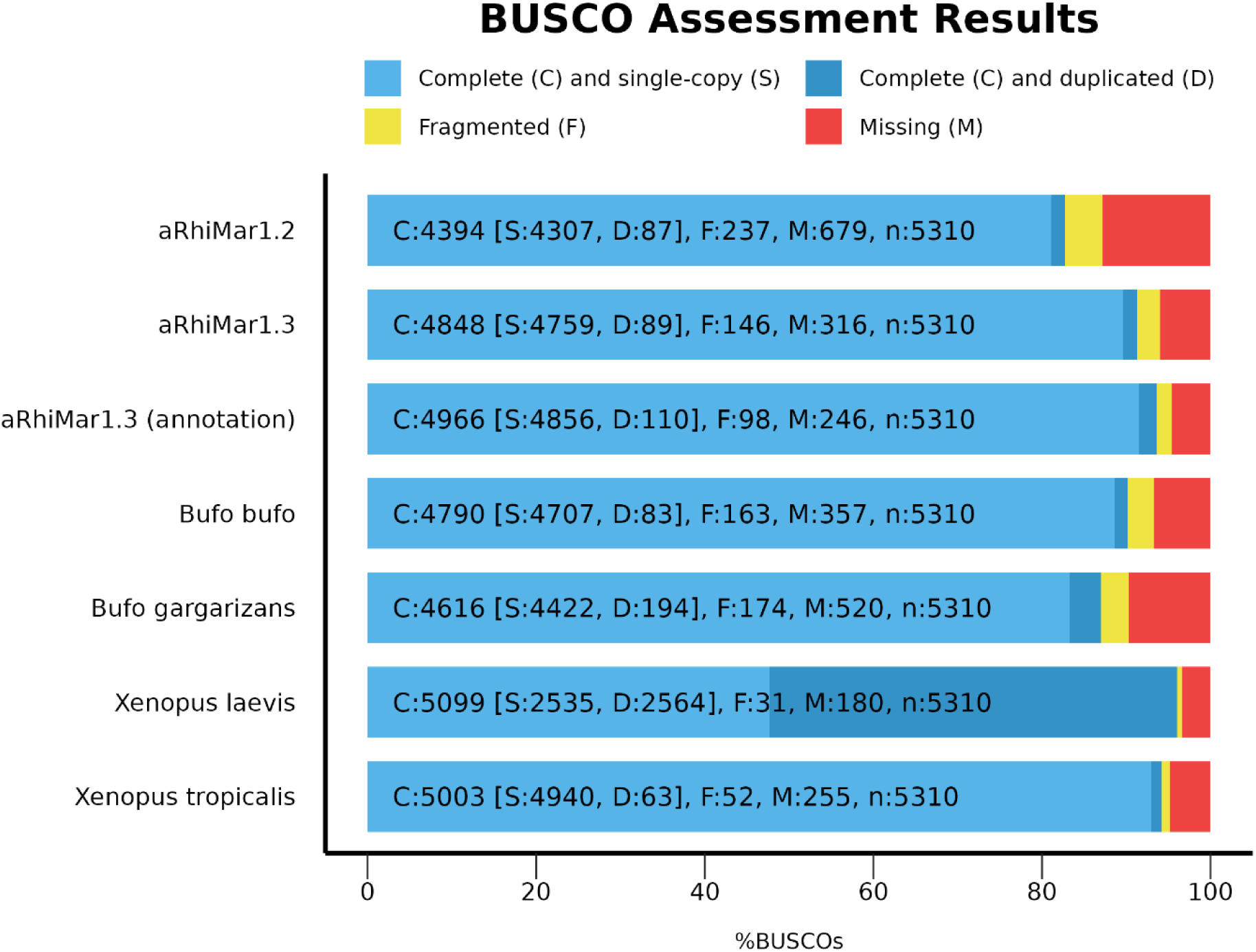
BUSCO assessment of the draft assembly (aRhiMar1.2), updated (aRhiMar1.3) cane toad (*Rhinella marina*) and four other amphibians genome assemblies.

For DepthKopy analysis (Figure 2), mean copy number of completed BUSCO genes in both versions were close to one, which indicated true single copies in the assembly. In aRhiMar1.2, copy number of duplicated BUSCO genes showed a bimodal distribution with peaks at around 0.5 CN and 1 CN, which indicated false duplication in the assembly. There were 709 (2.25%) and 223 (0.71%) sequences having a copy number greater than 2 and equal to 0, respectively, suggesting possible collapsed repeats and low-quality sequences. In aRhiMar1.3, the bimodal distribution in “Duplicated” was skewed towards 1 CN, suggesting that fewer false duplications appeared in the assembly than in the draft assembly. The copy number analysis of the 100 kbps window scan showed a mean value closer to 1 and fewer windows having CN greater than 1, suggesting that most collapsed repeats were also resolved through the addition of ONT reads. There is still a peak of contigs/scaffolds (Figure 2 “Sequences”) with a CN of approximately 0.5, indicating a number of false duplicates or haplotype-specific sequences. These could be the consequence of conservative duplicate removal with Diploidocus but could also represent some sequences specific to sex-chromosomes, because a heterogametic ZW individual was sequenced.

**Figure 2:**
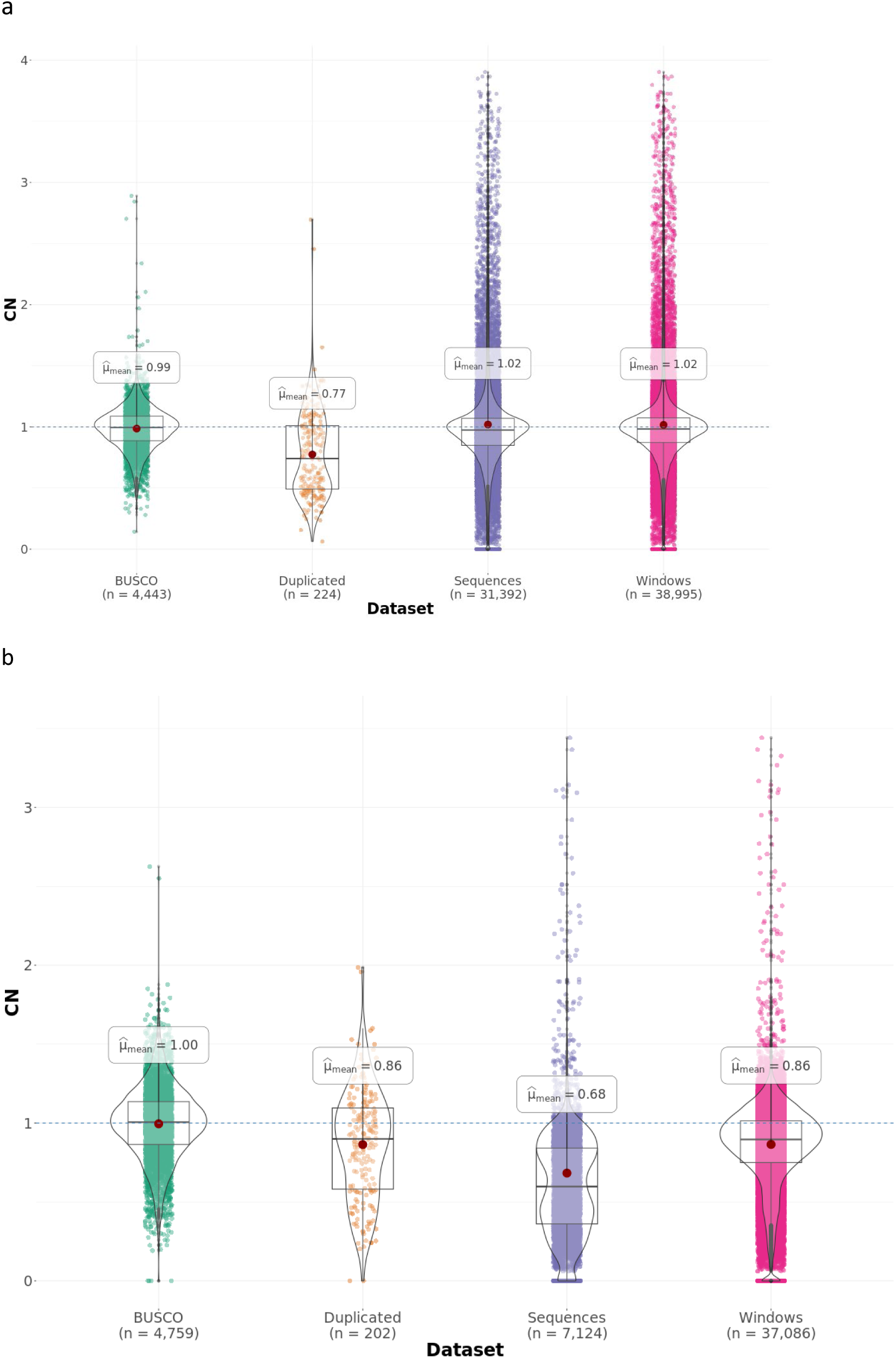
Copy number analysis of (a) the original *Rhinella marina* draft genome assembly (aRhiMar1.2) and (b) the updated genome assembly (aRhiMar1.3) based on the BUSCO results by DepthKopy. BUSCO: single-copy BUSCO genes; Duplicated: duplicated BUSCO genes; Sequences: contigs or scaffolds; Windows: 100,000 bps window.

### Nuclear genome annotation and evaluation

A total of 39,121 protein-coding genes with a median length of 2,535 bp were predicted in the cane toad genome assembly by GeMoMa across 2,879 scaffolds. BUSCO assessment of the longest protein per gene from the predicted proteome with 5,310 orthologous proteins in tetrapoda classified 93.6% as “Complete”, in which 2.1% were “Duplicated”. Nearly 5% of the proteins were “Missing” in the annotation (Figure 1).

Repeat annotation revealed that 72.48% of the genome sequences are repetitive elements (Table 2). More than half of the repeats were unclassified repetitive elements (42.29%). Class II transposable elements (TEs) (DNA transposons) were the most abundant repeat class (17.87%), dominated by hobo-Activator (13.16%). Class I TEs (retroelements) were only half of the class II TEs (9.29%), dominated by long interspersed nuclear elements (LINEs). A total of 1,338 high-confidence tRNAs were predicted.

**Table 2:** Repeat element summary of the draft (aRhiMar1.2) and the updated *Rhinella marina* genome assembly, aRhiMar1.3.

| **Repeats** | **aRhiMar1.2 (Mb)** | **aRhiMar1.3\* (Mb)** |
| --- | --- | --- |
| Class I TEs - Retroelements | 247.31 | 322.56 |
| Class II TEs - DNA transposons | 507.37 | 620.75 |
| Rolling circles | NA | 0.22 |
| Unclassified | 826 | 1,468.84 |
| Small RNA | 14.69 | 77.9 |
| Satellites | 11.3 | 4.39 |
| Simple repeats | 55.1 | 20.49 |
| Low complexity | 7.3 | 2.42 |
| **Total** | **1.63 Gb** | **2.51 Gb** |

\*Statistics included mtDNA

### NUMTs detection and validation in the cane toad

We chose three different approaches to detect NUMTs in the cane toad genome. Using NUMTFinder, a total of 64 putative NUMTs were found in the aRhiMar1.3 (Table 3). These were identified in 63 different scaffolds and all but one hit aligned to the control region of the mitogenome (Supplementary Table 2). Hit lengths ranged from 38 to 141 base pairs. There was one hit on scaffold 1637 that was 114 base pairs long, which corresponded to a partial *COX1* gene. When the assembly was masked with all default repeats, a total of 8 putative NUMTs remained detected where one come from a partial *COX1* gene and the remaining seven come from the control region (Supplementary Table 3). After visual examination of the sequence composition of the putative NUMTs (Supplementary Figure 2), the seven hits from the control region had clusters of AT microsatellite and extremely low GC content (0 – 7.14%), indicating these sequences were low-complexity regions of the genome assembly. Therefore, we decided to consider these seven hits from the control region as artefactual hits, along with all the control region hits from the unmasked genome.

**Table 3:** NUMTFinder results on the draft (aRhiMar1.2) and updated *Rhinella marina* genome assembly, aRhiMar1.3.

|  | **aRhiMar1.2** | **aRhiMar1.3** |
| --- | --- | --- |
| Genome size (Gb) | 2.55 | 3.47 |
| No. of NUMTs<br>(before quality control) | 42 | 64 |
| No. of NUMTs<br>(After quality control – default repeat-masking<br>(interspersed repeats, low complexity DNA)) | 0 | 1 |

Two methods of raw sequencing read analysis to detect NUMTs yielded different results. No NUMTs were detected using *dinumt*. Our analysis using *PALMER* with PacBio and Nanopore sequencing reads yielded non-overlapping results from the different sequencing technologies (Supplementary Table 4). In total, 49 (PacBio) and 45 (Nanopore) putative NUMTs were detected, each on a different scaffold. No putative NUMT was detected in both PacBio and Nanopore data sets. None of the 94 putative NUMTs detected by *PALMER* showed any supporting reads carrying NUMT insertion when visualised in IGV, with all the corresponding regions of aRhiMar1.3 well-supported by numerous spanning reads. Each putative NUMT read identified by *PALMER* mapped to a different part of the aRhiMar1.3 assembly, indicating that the putative NUMTs were likely to be the result of chimeric reads generated by library preparation or sequencing steps. All *PALMER* NUMT candidates were, therefore, excluded as false-positive signals.

### NUMTs detection in amphibian

In order to gain insight about the NUMT landscape across amphibians, we ran NUMTFinder on 16 amphibian genomes. Predicted NUMT counts across amphibians ranged from zero (*Xenopus tropicalis*) to 20,496 (*Leptobrachium leishanense*) (Table 4). We observed similar patterns as in the cane toad genome assemblies, with an abundance of putative NUMTs matching low-complexity control regions. Therefore, we adopted the same filtering regime used in the cane toad, and removed putative NUMTs corresponding to short hits to the mitogenome control region. This reduced the number of putative NUMTs by two orders of magnitude (0 – 356 NUMTs). Several species showed minimal reduction whilst, in an extreme case, all 20,496 putative NUMTs of *Leptobrachium leishanense* were filtered. NUMT fragments spanned across the mtDNA genome, from tRNA to protein-coding genes and rRNA (Supplementary Table 5). Visual inspection of the distribution of NUMT numbers after filtering revealed a dominant cluster of eleven species (65%, including cane toad) with 0-4 NUMTs, and the remaining six (35%) with 10+. To test the previous relationship of genome size and NUMT count, we therefore divided species into Low (<5) and High (≥10) NUMT counts. There was a significant genome size difference between the Low and High genomes after filtering (Wilcoxon test, *p* = 0.007), which was not seen with the unfiltered NUMTFinder results (Figure 3).

**Figure 3:**
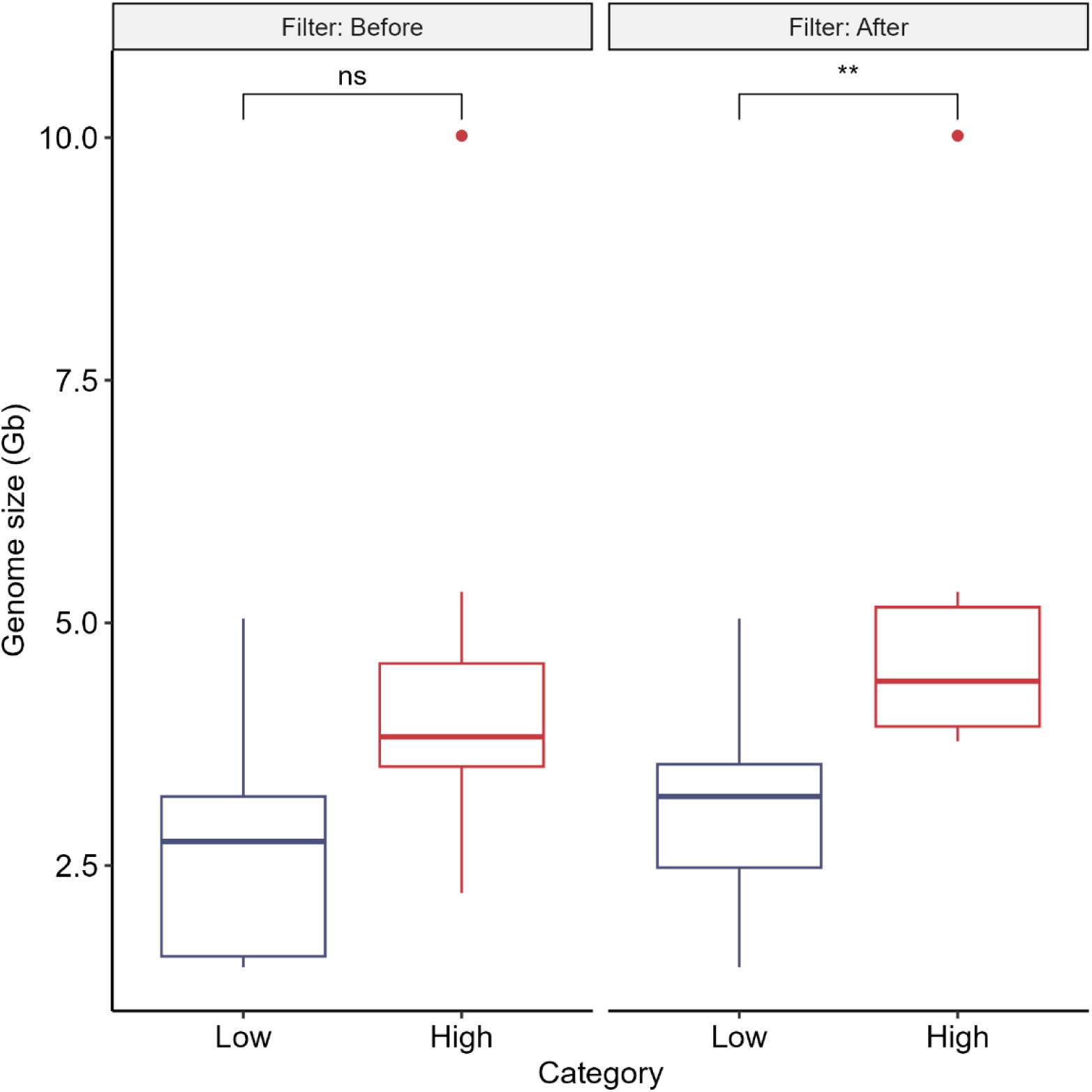
Relationship between genome size and the number of NUMT of 16 other amphibians and *Rhinella marina* found by NUMTFinder before and after custom filtering. Species with < 5 NUMTs are categorised as “Low” and those with ≥ 10 NUMTs as “High”. Statistical significance: ns indicates not significant and ** indicates p ≤ 0.001.

**Table 4:** NUMTFinder (BLASTN) results for 16 other amphibians and *Rhinella marina* (bottom) before and after filtering.

| Species | Genome accession number | mtDNA accession number | References | Genome Size (Gb) | No. of NUMTs | No. of NUMTs (After filtering) |
| --- | --- | --- | --- | --- | --- | --- |
| *Bombina bombina* | GCA_027579735.1 | NC_042501.1 | (Pabijan, et al. 2008; Rhie, et al. 2021) | 10.02 | 124 | 123 |
| *Bufo bufo* | GCA_905171765.1 | LR991678.1 | (Streicher, et al. 2021b) | 5.04 | 1 | 1 |
| *Bufo gargarizans* | GCA_014858855.1 | NC_008410.1 | (Cao, et al. 2006; Lu, et al. 2021) | 4.55 | 10 | 4 |
| *Dendropsophus ebraccatus* | GCA_027789765.1 | CM051072.1 | (Rhie, et al. 2021) | 2.21 | 87 | 0 |
| *Discoglossus pictus* | GCA_027410445.1 | CM050223.1 | (Rhie, et al. 2021) | 3.87 | 356 | 112 |
| *Geotrypetes seraphini* | GCA_902459505.2 | NC_020155.1 | (Ovchinnikov, et al. 2023; San Mauro, et al. 2009) | 3.78 | 36 | 33 |
| *Hymenochirus boettgeri* | GCA_019447015.1 | NC_015615.1 | (Bredeson, et al. 2024; Irisarri, et al. 2011) | 3.21 | 5 | 1 |
| *Leptobrachium ailaonicum* | GCA_018994145.1 | MZ394043.1 | (Li, et al. 2019b) | 3.54 | 170 | 2 |
| *Leptobrachium leishanense* | GCA_009667805.1 | NC_031411.1 | (Li, et al. 2019a; Liang, et al. 2016) | 3.55 | 20496 | 0 |
| *Microcaecilia unicolor* | GCA_901765095.2 | NC_023515.1 | (Ovchinnikov, et al. 2023; San Mauro, et al. 2014) | 4.69 | 34 | 33 |
| *Pyxicephalus adspersus* | GCA_004786255.1 | NC_044480.1 | (Cai, et al. 2019; Denton, et al. 2018) | 1.56 | 1 | 1 |
| *Rana temporaria* | GCA_905171775.1 | NC_042226.1 | (Chen 2018; Streicher, et al. 2021a) | 4.11 | 27 | 26 |
| *Rhinatrema bivittatum* | GCA_901001135.2 | NC_006303.1 | (Ovchinnikov, et al. 2023; San Mauro, et al. 2004) | 5.32 | 15 | 14 |
| *Xenopus borealis* | GCA_024363595.1 | NC_018776.1 | (Evans, et al. 2022; Lloyd, et al. 2012) | 2.75 | 2 | 1 |
| *Xenopus laevis* | GCA_017654675.1 | NC_001573.1 | (Roe, et al. 1985; Session, et al. 2016) | 2.74 | 19 | 1 |
| *Xenopus tropicalis* | GCA_000004195.4 | NC_006839.1 | (Bredeson, et al. 2024) | 1.45 | 0 | 0 |
| *Rhinella marina* |  | NC_066225.1 | This paper, (Cheung, et al. 2024) | 3.47 | 64 | 1 |

## Discussion

The presence of NUMTs can significantly confound results of mtDNA analyses. In the cane toad mitochondrial population genomics study, we did not identify any NUMT insertions (Cheung, et al. 2024). However, the results may have been limited by a relatively incomplete draft genome (aRhiMar1.2) and the use of only one detection method (NUMTFinder). To address these potential limitations, we improved the nuclear genome assembly and incorporated two additional NUMT detection methods (*dinumt* and *PALMER*). These new data suggested one newly found NUMT in the cane toad genome. Nevertheless, this NUMT appears to be old and divergent, and did not contribute to the haplotypes identified in the invasive cane toad population. Furthermore, we investigated how NUMT detection was affected by repeat content. In several amphibian genomes with high repeat content (no repeat-masking), we discovered that the number of NUMT was highly likely to be inflated due to the low complexity regions of the genome. These findings highlight the importance of accounting for repeat regions within the mtDNA when using a homology search approach.

### Revision to cane toad nuclear reference genome

Due to its repetitive nature, aRhiMar1.2 (Edwards, et al. 2018) appears to be highly collapsed, resulting in significant discrepancies in genome size estimates obtained from flow cytometry and densitometry (2.55 Gb vs 3.98 - 5.65 Gb). To address this issue, we assembled a more contiguous reference genome by leveraging multiple sequencing technologies, including PacBio long reads, Nanopore long reads, linked-read sets (10X Genomics) and short-read Illumina sequences. This combined data set yielded a 3.47 Gb haploid genome assembly, representing a substantial 0.9 Gb increase in genome size compared to aRhiMar1.2. This addition in the improved assembly coincides with the identification of an additional 0.88 Gb of repeats (Unclassified: 643 Mb; TEs: 189 Mb; Small RNA: 63 Mb) (Table 2), suggesting that a large portion of collapsed repeats has been resolved. This observation is supported by DepthKopy analysis, which indicated a reduction in collapsed repeats across different scaffolds (Figure 2). In addition, the contiguous assembly has enhanced the discovery and identification of repeat elements boundaries and relationships (e.g. fewer satellites, simple and low-complexity repeats). The assembly size fell within DepthSizer estimation range (3.4 – 3.5 Gb) and was much closer to the estimates obtained from flow cytometry and densitometry. It is notable that the margin of error in genome size estimation by flow cytometry could be significant, and we have previously found it to overestimate genome sizes in other species (Chen, et al. 2022a; Chen, et al. 2022b).

Beyond contiguity, the completeness and quality of the assembly also improved significantly with BUSCO completeness reaching 91.3%. This marks a substantial increase from 85.8% of aRhiMar1.2 with a 5.9% increase in single-copy BUSCO genes and a 3.5% decrease in missing BUSCO genes (Figure 1). This indicated that the additional ONT sequencing reads and use of polishing software have facilitated the assembly of more conserved protein orthologs (*n* = 452). Despite the assembly of the improved cane toad genome still being at the scaffold level, the BUSCO score is already comparable to other amphibian genome assemblies at the chromosome level. Specifically, the mean BUSCO of chromosomal-level assembly stands at 87.45 ± 0.09 % while scaffold-level assembly averages 81.56 ± 0.17 % (Zuo, et al. 2023). The relatively low BUSCO completeness of amphibian genomes could reflect particular challenges with assembling these repeat-rich species, or it could reflect taxonomic differences in gene content that are not currently captured by the BUSCO datasets. Similarly, the majority of Duplicated BUSCO genes in aRhiMar1.3 appear to be genuine duplicates based on read depth (Figure 2b).

### NUMT detection

Our previous mitochondrial study of the cane toad carefully examined the validity of the nuclear-derived sequencing reads for constructing the mitochondrial genome (Cheung, et al. 2024). Here, we aimed to clarify the NUMT landscape by producing an updated genome assembly with higher contiguity and completeness. Using the same method (NUMTFinder) as was used in Cheung et al. (2024), where no NUMTs were detected, with the improved genome we detected 64 putative NUMTs. After repeat-masking, only one from *COX1* gene remained suggesting that mostly these putative NUMTs from the control region of the mtDNA were highly likely to be false-positives. None of the haplotypes identified in introduced populations arose from this *COX1* NUMT. To make sure that the homology search approach was not missing NUMTs, we employed two alternative NUMT detection methods, *dinumt* and *PALMER*. These methods directly search for signatures of NUMT insertions, utilising raw sequencing reads. These two methods yielded consistent null results, suggesting the absence of NUMT in the cane toad nuclear genome. Of most importance, there is no evidence to suggest the presence of recent (high identity) NUMT fragments that might confuse analysis of real mtDNA sequencing. This finding further supports our previous mtDNA analyses of the cane toad and indicates that the new haplotypes identified in introduced populations are likely to be genuine, rather than NUMTs.

### Accounting for repeats in homology search

Repeat sequences pose challenges for homology-based searches. This issue was previously recognised and masking repetitive and low complexity regions in the nuclear genome prior to NUMT detection has been recommended to address this problem (Kielbasa, et al. (2011); Tsuji, et al. (2012). However, over-zealous masking of repeats could result in missing real NUMTs and may not be necessary where the mtDNA lacks repetitive regions. Amphibian genomes are known to be repeat-rich, ranging from 33.69% to 74.91% in the nuclear genomes (Zuo, et al. 2023) and 50% of amphibians have repeats in the control region of the mitochondrial genome (Formenti, et al. 2021). So, although care must be taken when masking repeats, it is also expected that random alignment to repeat-rich sequences will be observed when using NUMTFinder or similar techniques. Here, we demonstrate that performing the NUMT search with and without repeat-masking can provide a more nuanced view of the impact of repeats in homology-based NUMT detection.

We extrapolate our experience in detecting NUMT in the cane toad nuclear genome to other amphibian nuclear genomes. Among the 16 amphibian genomes tested, we observed that 10 of them (62.5%) had putative NUMTs originating from the control region of the mitochondrial genome. This result suggests that false-positive signals of NUMTs are common in repeat-rich nuclear genomes, rather than being specific to the cane toad genome. Based on our observations in a taxonomic group with high-repeat nuclear genome, we advise future research to carefully inspect the results from BLASTN (the most common way to detect NUMTs) or apply multiple methods to validate results. It is technically challenging to distinguish between genuine NUMTs transferred from the control region of the mitochondrial genome and false-positive NUMTs generated by the homology-based search software. For this work, we operate on the assumption genuine NUMTs will also include regions outside the repeat regions. Nevertheless, there is the risk that genuine NUMTs originating from the control region may inadvertently be removed by repeat masking, and the strategy employed should consider the relative risks of false positives and false negatives. Considering the tradeoff between accuracy and feasibility, more research should focus on developing alternative and less computation-intensive software.

### NUMT observation in amphibian

There has not been a genome-wide NUMT analysis in class Amphibia to date. The majority of the studies in amphibian have focused on a single mitochondrial gene, such as cytochrome b or cytochrome oxidase I, because of the utility of these genes in phylogenetics and phylogeography (Meng, et al. 2014; Vences, et al. 2014). Here, we found that the number of NUMTs across amphibian varies, ranging from none to a few hundred, similar to what has been found in avian genomes (Baltazar-Soares, et al. 2023; Liang, et al. 2018). In general agreement with previous work (Hazkani-Covo, et al. 2010), we found a weak positive relationship between genome size and the NUMT content. However, it was clearly non-linear and the majority of amphibian genomes had very few NUMTs regardless of genome size (Table 4). These findings suggested each species and taxon should be investigated independently to understand more about the NUMT evolution across the tree of life.

For *Xenopus tropicalis*, the only amphibian with previously documented NUMTs (Hazkani-Covo, et al. 2010), we found no NUMTs. The lack of details about how these NUMTs were identified hinders our ability to explain these divergent results. The previous analysis might have been looking for more ancient/degraded NUMTs, while in our study, we used different methodology and identified more recently transferred NUMTs. Also, different versions of *Xenopus tropicalis* genome assemblies (v4.0 or earlier vs v10.0) were used across studies. As expected for a more completed assembly, more NUMTs may be recovered from contigs breakpoints. It could also be that the previously identified NUMTs were assembly artefacts and were resolved using the updated assembly.

It is worth noting that the genome-wide NUMT search in this study, similar to that in avian genomes (Liang, et al. 2018), only considers one individual of each species for the presence/absence of NUMTs. Large-scale human studies have already shown that the composition of the NUMT landscape varies across different individuals (Wei, et al. 2022). In order to understand more about NUMTs in non-model species, multiple individuals should be included in future studies.

## Conclusion

To validate the absence of NUMTs in the cane toad nuclear genome, we significantly improved the genome assembly in terms of contiguity and completeness. We found one NUMT in this assembly across three different NUMT prediction methods. This NUMT was short and too divergent from the mitogenome to contribute false haplotypes in population genetics studies. Additionally, we investigated the NUMT landscape in a range of amphibian species, supporting previous observations for a weak relationship between genome size and number/content of NUMTs. We show how repetitiveness of the nuclear genome may confound the number of NUMTs reported in some species, highlighting a need to be clear and consistent in how NUMTs are to be defined if comparison across studies is to be made. Future studies should check for potential false-positive results, a common problem given the nature of these methods.

## Data availability

The data underlying this article are available in the European Nucleotide Archive with the study accession PRJEB24695 and assembly accession GCA_900303285.2.

## Acknowledgements

We thank the Ramaciotti Centre for Genomics, University of New South Wales (UNSW) who performed the sequencing in this study. This research includes computations using the computational cluster Katana supported by Research Technology Services at UNSW Sydney. This study was supported by the Australian Research Council (grant LP180100721 to RJE, grant DP160102991 to RS and LAR), the UNSW Scientia Program (to LAR), and the UNSW Scientia PhD Scholarship (to KC).

