## Supplementary Figure for "Repeat-rich regions cause false positive detection of NUMTs: a case study in amphibians using an improved cane toad reference genome"

Supplementary Figure 1: BUSCO analysis on 16 anuran and the cane toad genome assemblies (aRhiMar1.2 and aRhiMar1.3).

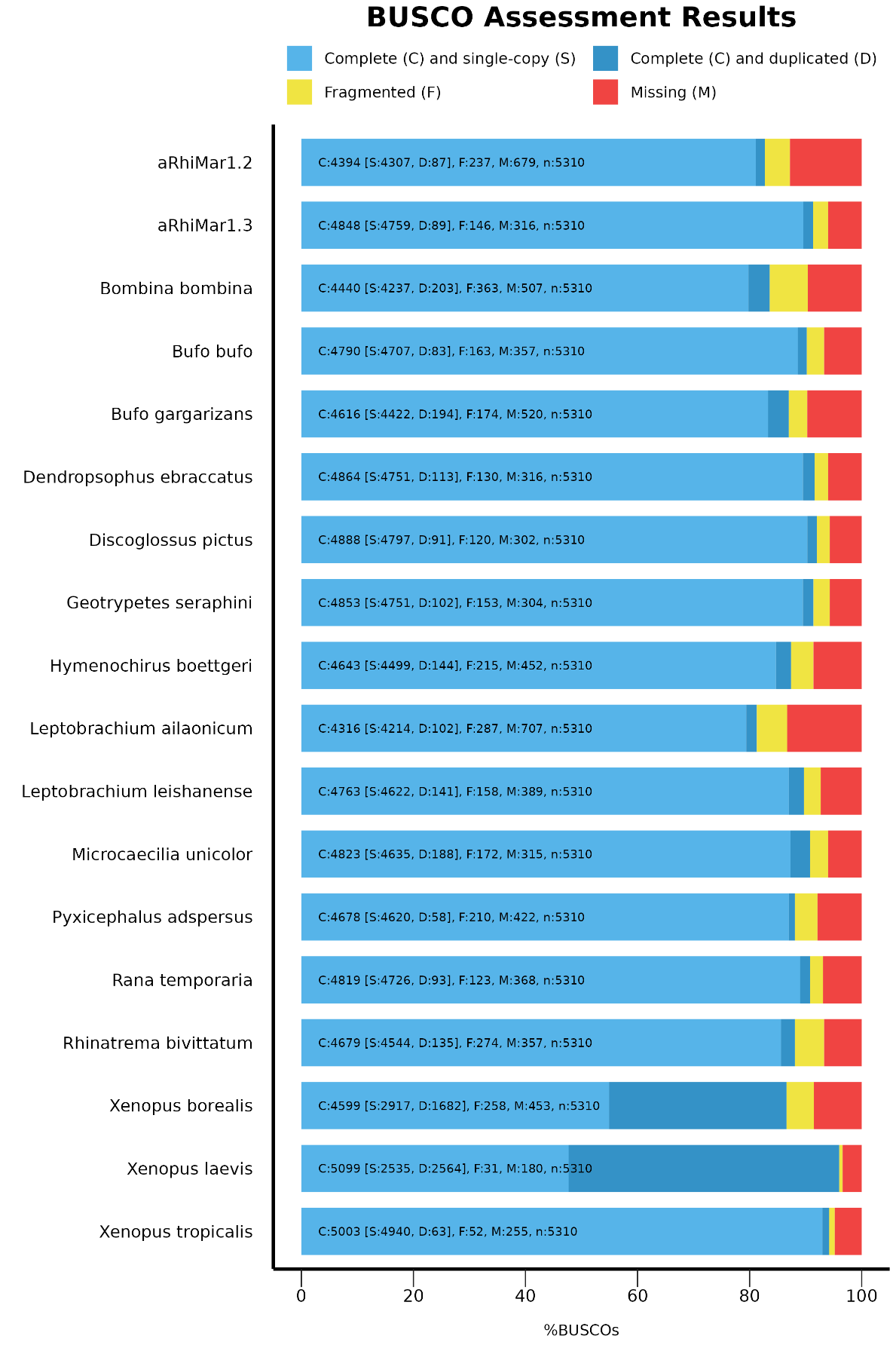

Supplementary Figure 2: Sequence composition of putative NUMTs found by NUMTFinder in aRhiMar1.3 after repeat-masking. Sequence contents are coloured based on the nucleotides (A: red, T: green, C: cyan, G: yellow). Except for *scaffold1637.253195-253308*, other putative NUMTs were rich in sequences with low-complexity.

>scaffold1637.253195-253308

GAGGAGAGGGGATCTGTAGAGTCATTCCACGTTAGTTGATGTTATGCTTGCAGAGATCAATAGACGTTTTGATGAGAAAGCTTCTTAAATAATAAATATTACATAATAATACAG

>scaffold1091.656950-657005

TAAAATTGAAAATTTTTATATATATATATATATATATATATATATATATATATTTT

>scaffold198.0314367-0314411

TTTAATATTCATATATATATATATATATATATATATATATAAAAA

>scaffold212.0538694-0538741

ATATATATATATATATATATATATATTAAATAACAATAAATAATAATA

>scaffold624.1337855-1337915

AATTTAAAATATATATATATATATATATATATATATATTATTTTTAAAAAAAAATTTTAAT

>scaffold718.3872772-3872816

ATATATATATATATATATATATATATTAAATGTAATAATTCTTAA

>scaffold86.3386787-3386831

TTTTTTTTTTATATATATATATATATATATATATATATTTTAAAT

>scaffold958.0965483-0965552

AAAAATATATATATATATATATATATATATATATATATATATTTTTTTTAAATTGGAAAAATCCCTTTAA
